# Cell-Hub Database: a consolidated multi-species ligand-receptor interaction resource for cell-cell communication analysis

**DOI:** 10.64898/2026.09.23.753685

**Authors:** Gaspard Macaux, Pascal Maire

**Author notes:** Corresponding author: Gaspard Macaux, 24 rue du Faubourg Saint-Jacques, 75014 Paris, France.

## Abstract

Cell-cell communication analysis by ligand-receptor interaction inference has become a standard component of single-cell transcriptomics workflows. However, existing ligand-receptor databases are fragmented across independent resources with heterogeneous formats, inconsistent gene nomenclature, and limited species coverage, requiring researchers to manually curate and convert interaction data before use. Here we present CellHub Database, a consolidated ligand-receptor interaction resource integrating 10 independently curated source databases (CellChatDB, CellPhoneDB, CellTalkDB, connectomeDB2025, NicheNet, NeuronChatDB, FlyPhoneDB2, cell2cell, PlantPhoneDB, and PlantCellChatDB) into 67,584 curated interactions across 21 species. To our knowledge, this is the first resource to provide CellChat-compatible ligand-receptor databases for Drosophila melanogaster and Caenorhabditis elegans, and the first to consolidate multiple plant ligand-receptor databases into a unified CellChat-native format covering five plant species. The curation pipeline standardizes gene symbol nomenclature per species, resolves cross-source duplicates using a priority system with full provenance tracking, and validates all gene symbols against reference databases. Users can load individual source databases or a pre-merged version, enabling flexible and transparent cell-cell communication analysis. CellHub Database is freely available as CellChat-native RDS files and integrated into Cell-Hub, an open-source R/Shiny application for no-code single-cell RNA sequencing analysis (companion paper).

## INTRODUCTION

Cell-cell communication inference in single-cell transcriptomics relies on curated ligand-receptor interaction databases. Tools such as CellChat^1^, CellPhoneDB^2^, and NicheNet^3^ each provide their own curated interaction lists, but these have been developed independently, with heterogeneous formats, gene nomenclature conventions, and species coverage, creating a fragmented landscape that complicates comprehensive analysis.

A researcher studying mouse cell-cell communication must currently choose between CellChatDB (∼3,300 interactions), CellTalkDB^4^ (∼2,000), or NicheNet (∼5,600), with no straightforward way to combine them or assess their overlap. The situation is more restrictive for non-mammalian organisms: No CellChat-compatible ligand-receptor database exists for Drosophila melanogaster or Caenorhabditis elegans, despite the availability of species-specific ligand-receptor resources for these organisms^5, 6^. For plant species, PlantCellChatDB^7^ provides CellChat-native interactions for five species, but no resource consolidates these with other plant databases such as PlantPhoneDB^8^ into a unified, cross-validated format.

Meta-resources such as OmniPath^9^ and LIANA+^10^ aggregate interaction data from multiple sources through programmatic interfaces, but do not provide pre-curated, tool-native files for direct use without format conversion, particularly for non-model organisms.

Here we present CellHub Database, a consolidated ligand-receptor resource that unifies 10 independently curated databases into 67,584 interactions across 21 species in CellChat-native format. A standardized curation pipeline ensures consistent gene nomenclature, cross-source deduplication with provenance tracking, and gene symbol validation against reference databases.

## DATABASE CONSTRUCTION

### Source databases

CellHub Database integrates 10 ligand-receptor interaction databases selected based on three criteria: availability of downloadable interaction data, evidence of manual or systematic curation, and compatibility with a pairwise ligand-receptor interaction format (Fig. 1).

**Figure 1:**
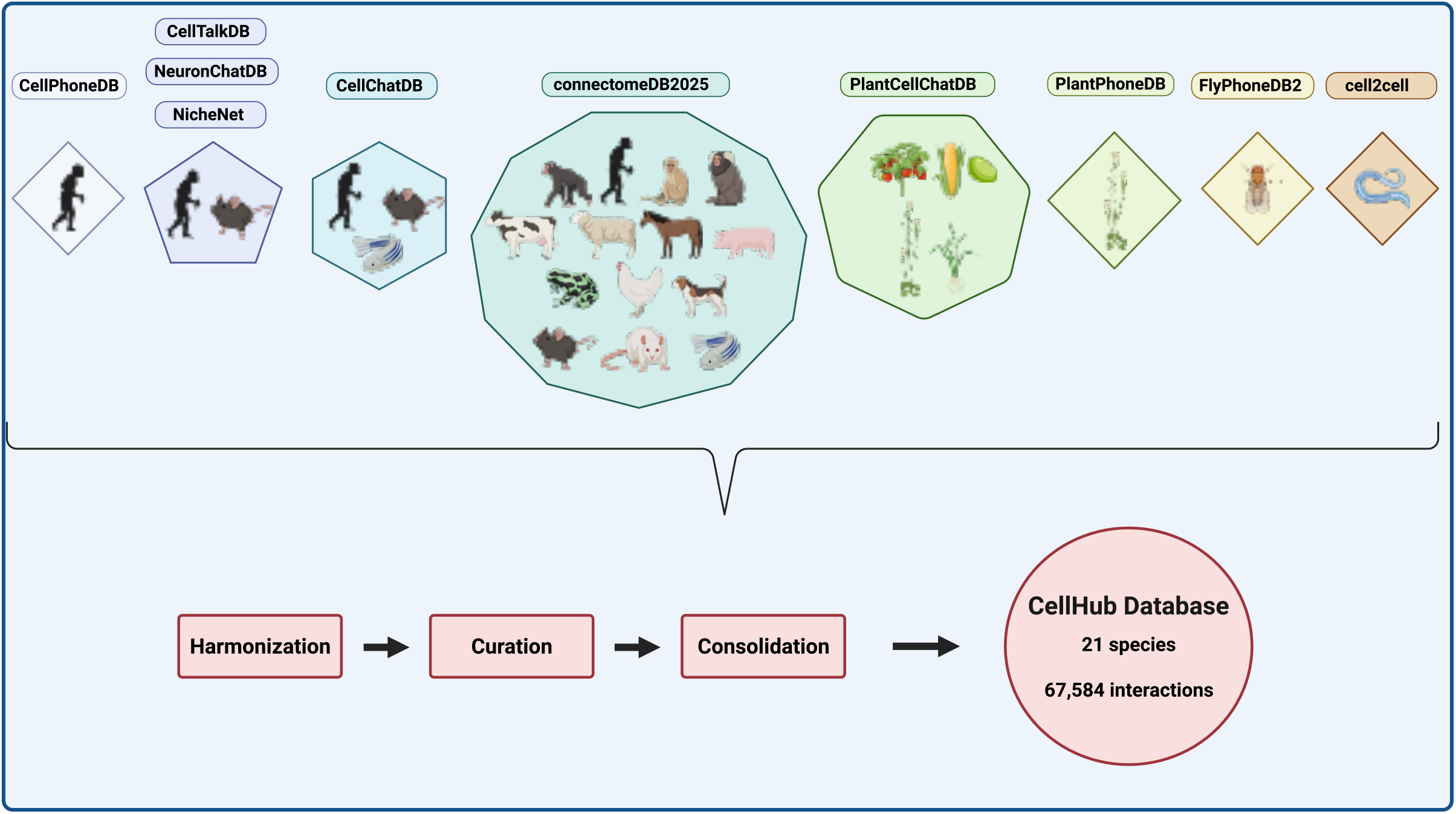
CellHub Database construction pipeline and species coverage. Ten ligand-receptor interaction databases are integrated through harmonization (format conversion, gene symbol normalization), curation (missing value handling, gene symbol validation), and consolidation (cross-source deduplication with provenance tracking) into 67,584 curated interactions across 21 species in CellChat-native format.

CellChatDB v2 is the reference database of the CellChat framework. It provides 3,233 human, 3,379 mouse, and zebrafish interactions manually curated from KEGG signaling pathway maps, with detailed pathway and interaction type annotations (Secreted Signaling, Cell-Cell Contact, ECM-Receptor). Interactions include multi-subunit receptor complexes defined in an accompanying complex table.

CellPhoneDB v5 provides 2,911 human interactions curated from literature and public protein databases. Partner identifiers use UniProt accessions rather than gene symbols, including 362 multi-subunit protein complexes. All entries were resolved to HGNC gene symbols using the database’s own gene and complex mapping tables during conversion.

CellTalkDB contributes 3,398 human and 2,033 mouse interactions manually curated from published literature. Each interaction is supported by one or more PubMed identifiers, providing direct bibliographic traceability.

connectomeDB2025^11^ is the largest single source, providing ligand-receptor pairs for 14 vertebrate species (human, mouse, rat, chimpanzee, macaque, marmoset, cow, dog, horse, pig, sheep, chicken, frog, zebrafish) with approximately 3,200 interactions per species. Interactions are classified as “Direct” (experimentally validated) or “Inferred” (mapped by cross-species orthology from human or mouse reference pairs).

The NicheNet v2 ligand-receptor network aggregates 5,098 human and 5,681 mouse interactions compiled computationally from multiple public resources including KEGG, Reactome, and literature-mined databases. Each interaction carries a resource count and directionality annotation.

NeuronChatDB^12^ provides 190 human and 183 mouse neuro-specific interactions covering five signaling categories: neurotransmitters (109 mouse), neuropeptides (37), gap junction proteins (20), synaptic adhesion molecules (9), and gaseous signals (8). Neurotransmitter and gas interactions use metabolite-enzyme composites (e.g. “GABA-GAD1”) to represent non-protein ligands through their biosynthetic enzymes. During conversion, these composites were expanded into individual enzyme-receptor pairs (e.g. Gad1-Slc32a1, Gad2-Slc32a1), increasing the mouse count from 183 to 452 interactions, using the enzyme as an expression proxy following the approach used internally by NeuronChat.

FlyPhoneDB2 is the only curated ligand-receptor database for Drosophila melanogaster, providing 1,798 interactions at three confidence levels: high (310), moderate (651), and low (837). Gene symbols follow FlyBase conventions (e.g. dpp, Toll, sNPF).

The cell2cell compendium provides 245 manually curated interactions for Caenorhabditis elegans across 20 functional categories including TGF-B signaling, Wnt signaling, cell adhesion, and neuronal communication. Confidence levels distinguish literature-supported (181), putative based on data (21), and putative based on literature (43) interactions. Gene symbols follow WormBase conventions (e.g. daf-7, sma-6).

PlantPhoneDB contributes 3,514 Arabidopsis thaliana interactions derived from seven resources (STRING, IntAct, BioGRID, interactome2.0, plant.MAP, literature, and orthologs). Gene identifiers use TAIR AGI locus IDs (e.g. AT1G01900).

PlantCellChatDB extends plant coverage to five species: Arabidopsis (5,481 interactions), maize (2,316), rice (1,864), soybean (1,457), and tomato (1,069), provided in CellChat-native format with pathway annotations.

### Harmonization

After format conversion, all interactions undergo harmonization to ensure cross-source compatibility.

Interaction type standardization. Variant spellings and source-specific classifications are normalized to four canonical categories: Secreted Signaling, Cell-Cell Contact, ECM-Receptor, and Non-protein Signaling. For example, NeuronChatDB’s “Neuropeptide” is mapped to Secreted Signaling, and “Gap junction protein” to Cell-Cell Contact.

Gene symbol normalization. Gene symbols are normalized to species-appropriate conventions using geneInfo reference tables. Human symbols follow HGNC convention (all uppercase, e.g. TGFB1). Mouse symbols follow MGI convention (first letter uppercase, e.g. Tgfb1). Rat symbols follow RGD convention, identical to MGI (e.g. Tgfb1). The remaining 8 mammalian species (chimpanzee, macaque, marmoset, cow, dog, horse, pig, sheep) use HGNC uppercase convention as provided by connectomeDB2025. Zebrafish symbols follow ZFIN convention (all lowercase, e.g. tgfb1). Chicken symbols use uppercase convention (CGNC). Frog symbols follow Xenbase convention (all lowercase, e.g. tgfb1), as provided by connectomeDB2025. Drosophila symbols follow FlyBase convention, which preserves the original mixed casing as published (e.g. dpp, Toll, sNPF). C. elegans symbols follow WormBase convention (lowercase with hyphens, e.g. daf-7, sma-6). Arabidopsis identifiers use TAIR AGI locus IDs (e.g. AT1G01900). The remaining four plant species (maize, rice, soybean, tomato) use the identifiers provided by PlantCellChatDB, which follow species-specific genome database conventions (e.g. OS07G0173700 for rice, SOLYC10G076410.1 for tomato).

Evidence normalization. Bibliographic references are standardized to a uniform PMID format (e.g. PMID:12345678). Free-text entries used as evidence and aberrant identifiers are cleaned or removed.

### Curation

Missing value normalization: Empty strings and whitespace-only values are replaced with NA across all columns for consistent missing-data handling.

Pathway name cleaning: Database names erroneously stored as pathway annotations (e.g. “MultiNiche”, “CellTalk”) are set to NA to prevent confusion between source provenance and biological pathway classification.

Annotation cleaning: Generic annotations carrying no biological information (e.g. “Signaling” without further specification) are removed.

UniProt ID removal: Some source databases contain residual UniProt accession numbers (e.g. Q9Y5Y7) in the ligand or receptor fields instead of gene symbols. These unresolvable identifiers are flagged and removed: 519 entries were excluded, primarily from connectomeDB2025 (284) and CellChatDB v2 (196).

Gene symbol validation: Each ligand and receptor is validated against geneInfo reference tables and CellChat complex name tables (case-insensitive matching). Metabolite-enzyme composite names (e.g. GABA-GAD1) are validated by checking that the enzyme component is a recognized gene symbol. Interactions containing unresolvable identifiers are removed and logged to an audit file: 185 entries were removed for invalid gene symbols, primarily pseudogenes and obsolete nomenclature from NicheNet (n=137) and connectomeDB2025 (n=37).

Interaction name reconstruction: After gene symbol normalization and validation, interaction identifiers are rebuilt from the normalized ligand and receptor symbols to ensure internal consistency (e.g. Tgfb1_Tgfbr1 for mouse, TGFB1_TGFBR1 for human).

### Consolidation

Interactions from all sources for a given species are pooled, and canonical ligand-receptor pair matching (case-insensitive) identifies duplicates. When the same pair exists in multiple databases, the entry from the highest-priority source determines the primary record, while metadata (evidence, pathway, interaction type) from lower-priority sources is merged. Priority reflects curation quality: CellChatDB v2 > connectomeDB2025 > CellPhoneDB > NeuronChatDB > PlantCellChatDB > cell2cell > PlantPhoneDB > FlyPhoneDB2 > CellTalkDB > NicheNet. A provenance column records all contributing databases for each interaction, enabling users to assess cross-source validation.

### Output format

The pipeline produces two types of output per species in CellChat-native RDS format. Individual source files (one per contributing database per species) preserve curated interactions from each source independently, enabling source-specific analyses and reproducibility. Merged files contain all interactions for a given species deduplicated across sources, providing maximum interaction coverage.

Each RDS file is a list containing four elements. The interaction table stores one ligand-receptor pair per row with ligand and receptor symbols, pathway name, interaction type, annotation, bibliographic evidence, winning source database, and a provenance field listing all contributing databases. The complex table defines multi-subunit protein complexes, mapping complex names to their individual gene components. The cofactor table lists cofactors and co-receptors associated with specific interactions. The geneInfo table provides gene symbol references for validation and expression matrix matching.

This format is directly compatible with CellChat’s createCellChat() function via assignment to the @DB slot, requiring no format conversion. The files can also be loaded in standalone R scripts as standard RDS objects using readRDS().

## DATABASE CONTENT

### Species coverage and interaction counts

CellHub Database comprises 67,584 species-specific curated interactions across 21 species, representing 21,882 unique biological ligand-receptor pairs when compared across species using case-insensitive gene symbol matching. The redundancy ratio of 3.1x reflects the contribution of connectomeDB2025, which maps approximately 3,200 conserved pairs across 14 vertebrate species by orthology (Fig. 2a).

**Figure 2:**
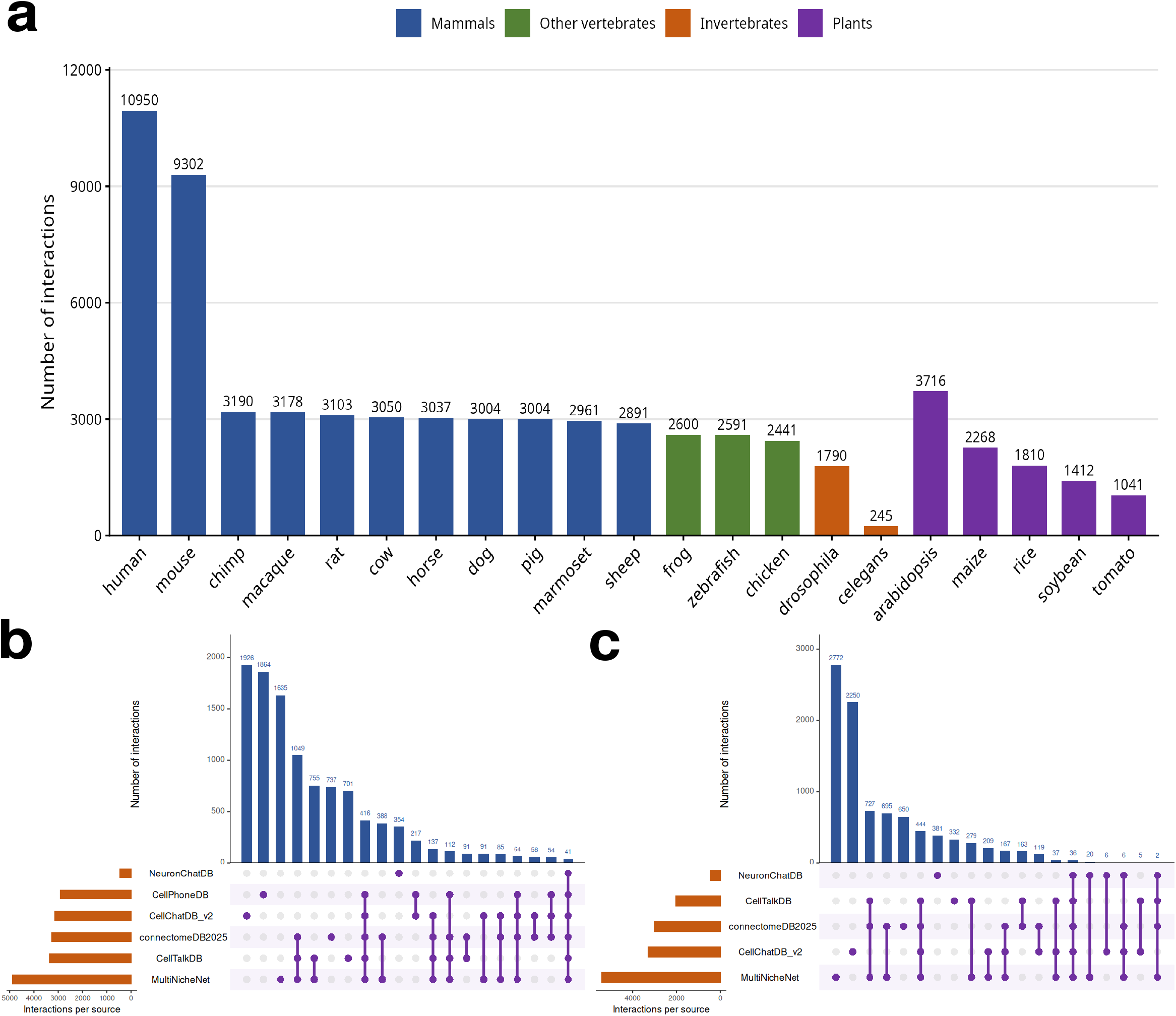
CellHub Database content and cross-source validation. (a) Number of curated interactions per species, colored by organism category. (b) UpSet plot showing the intersection of ligand-receptor pairs across 6 source databases for human (10,950 interactions). (c) UpSet plot for mouse (9,302 interactions from 5 sources). Vertical bars indicate the number of interactions in each source combination, horizontal bars indicate total interactions per source.

The database covers four organism categories: 11 mammals (human, mouse, rat, chimpanzee, macaque, marmoset, cow, dog, horse, pig, sheep), 3 other vertebrates (chicken, frog, zebrafish), 2 invertebrates (Drosophila melanogaster, Caenorhabditis elegans), and 5 plants (Arabidopsis thaliana, maize, rice, soybean, tomato). Human provides the highest coverage with 10,950 interactions from 6 sources, followed by mouse with 9,302 interactions from 5 sources. The remaining vertebrate species contain between 2,441 (chicken) and 3,190 (chimpanzee) interactions, each from a single source (connectomeDB2025). Among plants, Arabidopsis has the highest coverage with 3,716 interactions from two sources (PlantCellChatDB and PlantPhoneDB).

### Cross-source validation

For human and mouse, the availability of multiple independent source databases enables cross-source validation of individual interactions (Fig. 2b, c).

In human, 3,733 interactions (34.1%) are supported by at least two independent databases. Among these, 1,679 are confirmed by exactly 2 sources, 1,263 by 3 sources, 331 by 4 sources, 419 by 5 sources, and 41 interactions are present in all 6 databases simultaneously. The most frequent cross-source combination is connectomeDB2025, CellTalkDB, and NicheNet (1,049 shared pairs), reflecting the partial overlap between orthology-based, literature-curated, and computationally aggregated approaches.

In mouse, 2,917 interactions (31.4%) are multi-source. The overlap structure is similar, with connectomeDB2025, CellTalkDB, and NicheNet sharing 727 pairs, and connectomeDB2025 and NicheNet sharing 695 pairs.

Each source contributes both shared and exclusive interactions. For human, CellChatDB v2 provides 1,926 exclusive pairs (61.4% of its content), CellPhoneDB provides 1,864 exclusive pairs (64.1%), and NeuronChatDB provides 354 exclusive pairs (75.6%), the majority representing neurotransmitter and neuropeptide signaling not covered by other databases. Conversely, CellTalkDB has the lowest exclusivity (701 exclusive pairs, 20.8%), indicating that most of its content is independently curated in other resources, which can be interpreted as strong cross-validation of its interactions.

Species covered by a single source (all vertebrates except human, mouse, and zebrafish, as well as Drosophila, C. elegans, and individual plant species) lack cross-source validation by definition. For these species, interaction confidence relies on the curation quality of the contributing database and, for connectomeDB2025 vertebrates, on whether the interaction is classified as “Direct” (experimentally validated) or “Inferred” (mapped by orthology).

### Interaction type and pathway annotations

Interaction types are annotated for 53.2% of human interactions: Secreted Signaling (4,644, 42.4%), Cell-Cell Contact (820, 7.5%), Non-protein Signaling (360, 3.3%), and ECM-Receptor (6, 0.1%). The remaining 46.8% lack type annotation, primarily interactions from NicheNet, CellTalkDB, and connectomeDB2025, which do not provide this classification in their source data. For mouse, 36.7% of interactions carry type annotations with a similar distribution.

### Gene coverage

For human, CellHub Database covers 1,785 unique ligand identifiers, 1,711 unique receptor identifiers, and 2,689 unique individual gene symbols (after splitting multi-subunit complexes). 431 genes (16.0%) function as both ligand and receptor in different interaction contexts. For mouse, the database covers 1,707 ligands, 1,584 receptors, and 2,677 unique gene symbols, with 422 dual-role genes. Human-mouse ortholog comparison reveals 7,618 shared ligand-receptor pairs, representing 81.9% of mouse and 69.6% of human interactions, consistent with the known conservation of mammalian signaling pathways.

## AVAILABILITY AND USAGE

CellHub Database is distributed as a single RDS file containing all 21 species freely available on the Cell-Hub Database GitHub repository (https://github.com/GaspardMacaux/CellHub-Database) under the MIT license. No registration, login, or authentication is required.

Within Cell-Hub (companion paper), users select a species from 21 available organisms through a graphical interface, then choose between loading the complete merged database or selecting individual source databases via checkboxes (Fig. 3a, b). Upon loading, a summary panel displays the number of interactions, contributing sources, pathway coverage, and multi-source validation statistics (Fig. 3c). The database is then assigned to a CellChat object for ligand-receptor interaction inference and visualization (Fig. 3d). The resulting bubble plots (Fig. 3e) and chord diagrams (Fig. 3f) enable visualization of inferred ligand-receptor interactions and pathway-level communication networks between cell populations. This dual-access mode enables researchers to either maximize interaction coverage (merged mode) or restrict analysis to specific curated sources for reproducibility with a given reference database.

**Figure 3:**
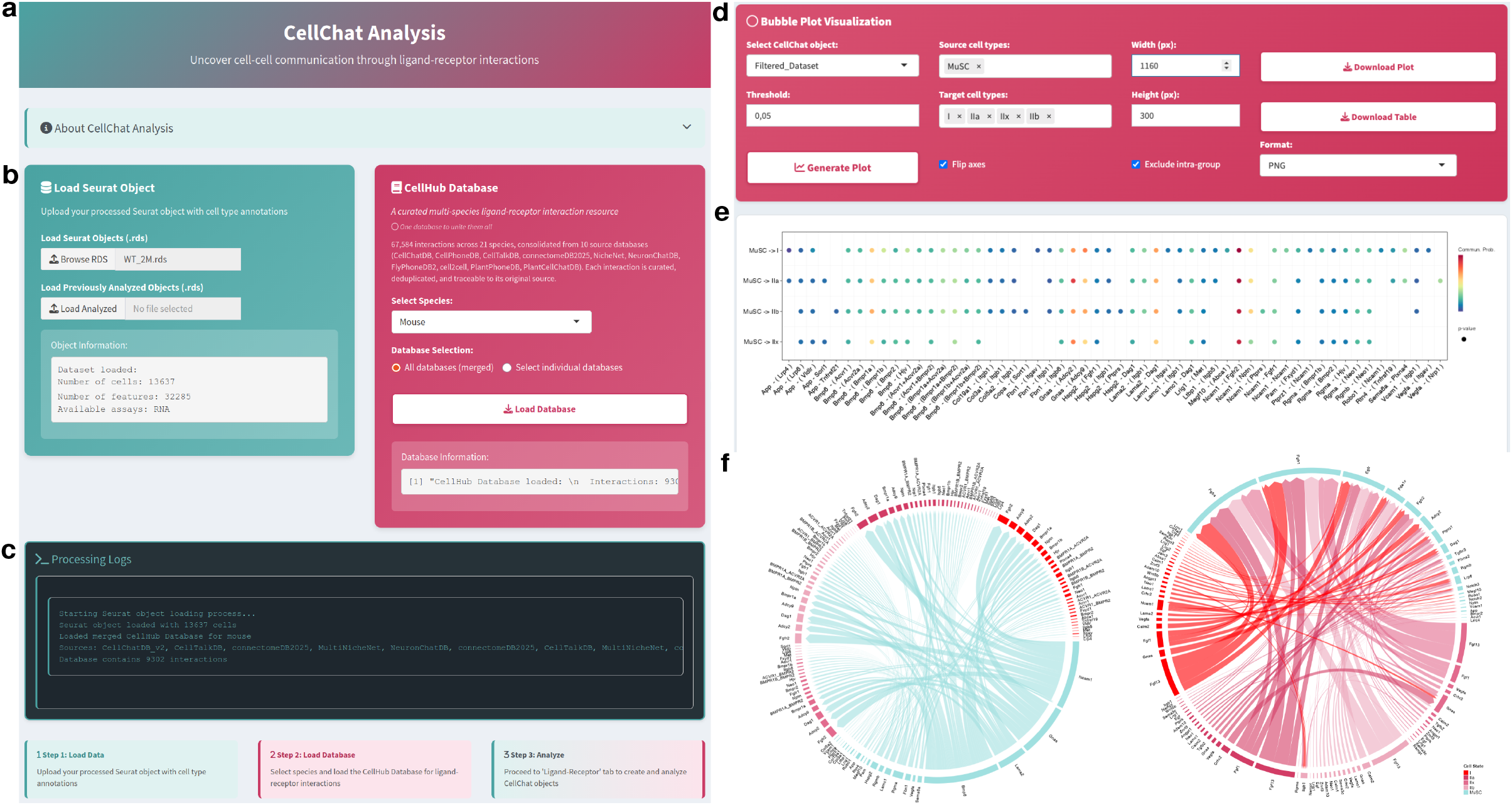
CellHub Database integration in Cell-Hub. (a) CellChat analysis module header. (b) Data loading interface showing Seurat object import (left) and CellHub Database selection panel (right) with species dropdown, database selection mode, and database information after loading (9,302 mouse interactions). (c) Processing logs confirming database loading with source provenance (CellChatDB_v2, CellTalkDB, connectomeDB2025, NicheNet, NeuronChatDB) and step-by-step workflow guide. (d) Bubble plot configuration interface with source and target cell type selection. (e) Bubble plot visualization of inferred ligand-receptor interactions between MuSC and myofiber subtypes, showing communication probability and significance. (f) Chord diagrams displaying cell-cell communication networks at pathway level.

For standalone use, the database can be loaded in any R session with a single command:

*db <- readRDS(“CellHubDB*.*rds”)*

The species of interest assigned to a CellChat object via:

*cellchat@DB <- db[[“mouse”]]*

Requiring no additional package installation beyond CellChat itself.

The complete curation pipeline, including all 10 conversion scripts, the harmonization and curation pipeline, quality control tools, and integrity verification scripts, is publicly available on the repository to ensure full reproducibility.

## DISCUSSION

CellHub Database addresses the fragmentation of ligand-receptor interaction resources by consolidating 10 independently curated databases into a unified, harmonized format directly usable by existing cell-cell communication tools. While aggregation frameworks such as LIANA+ and OmniPath provide programmatic access to multiple ligand-receptor resources, CellHub Database differs in scope by offering pre-curated, CellChat-native files for 21 species including non-model organisms, accessible through a graphical interface or a single R command without format conversion or additional dependencies. To our knowledge, it is the first resource to provide CellChat-compatible ligand-receptor databases for Drosophila melanogaster and Caenorhabditis elegans, and the first to consolidate multiple plant databases into a unified CellChat-native format covering five plant species.

The cross-source provenance tracking enables a new dimension of interaction assessment. For human and mouse, 34% and 31% of interactions respectively are confirmed by at least two independent sources, providing an orthogonal confidence metric beyond individual database evidence annotations.

Several limitations should be noted. CellHub Database currently uses CellChat-native format as its sole output; users of other tools such as CellPhoneDB or LIANA+ would need to convert the data to their respective input formats. Species covered by a single source lack cross-source validation, and their interaction coverage depends entirely on the contributing database. The gene symbol validation relies on CellChat’s geneInfo tables, which may not include all valid symbols for every species. Metabolite-mediated signaling is covered through NeuronChatDB’s enzyme-proxy approach for neurotransmitters and neuropeptides, but broader metabolite signaling (lipids, amino acids) remains underrepresented.

Future developments include integration of additional source databases such as NATMI and ICELLNET, expansion to additional species, and versioned releases synchronized with source database updates.

## Acknowledgements

We thank Edgar Jauliac and Léa Delivry for bioinformatics support, and Stéphanie Backer and Florian Britto for helpful comments on the manuscript.

## Ethics, Consent to Participate, and Consent to Publish declarations

All animal experiments were conducted according to the National and European legislation and institutional guidelines for the care and use of laboratory animals approved by the French government (Ministère de l’Enseignement Supérieur et de la Recherche, autorisation APAFiS #44987-2023092714211861 v4 approved on February 5, 2024, “Evaluation of gene therapy approaches and antisense strategies for genetic diseases”). No human participants were involved in this study; therefore, consent to participate and consent to publish are not applicable.

## AI assistance declaration

During the preparation of this manuscript, the authors used Claude (Anthropic) to assist with manuscript revision and to assist in portions of the Cell-Hub software development. All content was reviewed, verified, and approved by the authors, who take full responsibility for the integrity of the work.

## Data availability statement

Cell-Hub is freely available as a Docker image at Docker Hub (gaspardmacaux/cell-hub:latest) and the source code is available at https://github.com/GaspardMacaux/Cell-Hub. Cell-Hub Database is distributed within the Docker image and available on the GitHub repository https://github.com/GaspardMacaux/CellHub-Database. RNA sequencing data used for demonstration purposes have been deposited in the Gene Expression Omnibus (GEO) under accession number GSE309529 (https://www.ncbi.nlm.nih.gov/geo/query/acc.cgi?acc=GSE309529).

